# Robust genome editing via modRNA-based Cas9 or base editor in human pluripotent stem cells

**DOI:** 10.1101/2022.05.24.493220

**Authors:** Tahir Haideri, Alessandro Howells, Yuqian Jiang, Jian Yang, Xiaoping Bao, Xiaojun Lance Lian

**Affiliations:** Department of Biomedical Engineering, Pennsylvania State University, University Park, PA, 16802, USA; Department of Biology, Pennsylvania State University, University Park, PA, 16802, USA; The Huck Institutes of the Life Sciences, Pennsylvania State University, University Park, PA, 16802, USA; Davidson School of Chemical Engineering, Purdue University, West Lafayette, IN, 47907, USA

**Author notes:** Correspondence should be addressed to: Xiaoping Bao and Xiaojun Lance Lian. Co-first author (Tahir Haideri, Alessandro Howells).

## Abstract

CRISPR systems have revolutionized biomedical research because they offer an unprecedented opportunity to explore the application of genome editing in human pluripotent stem cells (hPSCs). Due to the inherent simplicity of CRISPR systems, requiring a Cas protein and its corresponding single guide RNA (sgRNA), they are more widely adopted and used for diverse biomedical research than their predecessors (zinc finger nucleases and TALENs). However, a bottleneck of applying CRISPR systems in hPSCs is how to deliver CRISPR effectors easily and efficiently into hPSCs. Herein, we developed modified mRNA (modRNA) based CRIPSR systems that utilized Cas9 or base editor (ABE8e) modRNA for genome editing of hPSCs via simple lipid-based transfection. We have achieved 71.09% ± 9.13% and 68.53% ± 3.81% gene knockout (KO) efficiency with Cas9 modRNA and ABE8e modRNA, respectively, which is significantly higher than plasmid-based systems. In summary, we demonstrate that our non-integrating modRNA based CRISPR methods hold great promise as the more efficient and accessible techniques for genome editing of hPSCs.

## INTRODUCTION

CRISPR-Cas systems are widely used for genome editing in a wide variety of cell types and are especially useful for high-throughput genome-wide screens(Xu et al., 2020; Yilmaz et al., 2018). Cas9 is the most used endonuclease of the CRISPR-Cas family(Cong et al., 2013; Mali et al., 2013) and can precisely cleave genomic DNA via double-stranded breaks (DSBs) when paired with a programmable single guide RNA (sgRNA) with minimal off-target effects. Repair of DSBs can occur through one of the two intrinsic pathways in mammalian cells: non-homologous end joining (NHEJ) and homology-directed repair (HDR). NHEJ results in insertions or deletions (indels) which can lead to frameshift mutations and consequently, gene knockout (KO)(Lian et al., 2016). Alternatively, co-delivery of a donor DNA template can precisely introduce desired sequence edits via the HDR pathway. DNA cleavage is mediated by the HNH and RuvC domains of the Cas9 protein(Cong et al., 2013; Mali et al., 2013). Mutations in these domains result in a catalytically inactive Cas9 (dCas9), which allows for a more general platform for RNA-guided, genomic delivery of a wide variety of covalently tethered effector proteins, among them being base editors(Richter et al., 2020). Two primary base editors used in practice are either adenosine or cytidine deaminases, also known as adenine base editors (ABEs)(Gaudelli et al., 2017; Richter et al., 2020) or cytidine base editors (CBEs)(Yu et al., 2020). ABEs perform specifically A to G conversion reactions while CBEs convert C to T. When covalently tethered to a dCas9, this enables researchers to introduce a genomic point mutation at high fidelity without DSBs, thus significantly reducing the risk of potentially detrimental indels and chromosomal rearrangements at off-target sites. ABEs and CBEs have been leveraged to correct disease-related point mutations and for gene KO purposes, at relatively high efficiencies and specificities(Antoniou et al., 2021; Kluesner et al., 2021).

Human pluripotent stem cells (hPSCs) can be expanded almost indefinitely while still maintaining their ability to differentiate into all somatic cell lineages(Jiang et al., 2021; Lian et al., 2012, 2013, 2014, 2015). They can be utilized to generate *in vitro* cell culture models for studying human development and disease modeling when coupled with CRISPR-Cas9 systems(Antoniou et al., 2021). Despite their remarkable potential for genome editing, the current state-of-the-art methods for delivering CRISPR components into hPSCs are far from ideal. Virus-mediated gene delivery is considered an efficient method for the delivery of CRISPR components into most cell types(Hsu et al., 2019). Commonly used viral vectors include lentiviruses, adeno-associated viruses (AAVs), and adenoviruses. Lentiviruses are normally integrating, which can increase the risk of tumorigenicity, and therefore, hPSC lines with lentiviral integrations may be counterproductive during their use in cell-based therapies. Additionally, hPSCs were reported to be resistant to lentiviral infection due to unique intrinsic immunity(Wu et al., 2018). AAVs and adenoviruses are two non-integrating alternatives to lentiviruses. However, adenoviruses are known to trigger high levels of innate immune response in transduced cells which can lead to inflammation, while AAVs have a relatively low packaging limit (∼4.7Kb), making it difficult to deliver CRISPR components. Additionally, AAVs and adenoviruses are laborious to produce and require the use of specialized equipment for their purification.

Non-viral methods include a variety of physical and chemical delivery strategies. Electroporation and lipid nanoparticles (LNPs) are two commonly used non-viral delivery methods, which use plasmid DNA for the delivery of CRISPR components via either transfection reagents or nucleofection(Liu et al., 2016). These methods however can be cytotoxic to cells and have lower transfection efficiency. Ribonucleoproteins (RNPs) consisting of Cas9 protein complexed with a sgRNA delivered via electroporation have also been shown to efficiently edit the genome(Martin et al., 2019). However, producing purified Cas9 protein can be cumbersome and not feasible for many labs.

An emerging alternative to these approaches is the use of chemically modified RNA (modRNA) for the delivery of CRISPR effectors into cells. The use of modRNA carries several advantages over the previous methods: 1) it is non-integrating, 2) it does not require transport across the nuclear membrane for expression (as is the case with plasmid DNA delivery), therefore increasing transfection efficiency, 3) it is relatively quick and easy to perform, 4) it requires a minimal starting cell population, and 5) it is only transiently expressed thus greatly reducing the risk of off-target activity.

In this study, we developed completely modRNA-based genome editing systems for hPSCs that utilize simple lipid-based transfection of sgRNAs, with Cas9 or ABE8e (base editor) modRNA. Using our optimized protocol, we were able to achieve up to 84% KO efficiency in hPSCs.

## RESULTS

### RNA based delivery of CIRSPR/Cas9 components can successfully knock out genes in hPSCs

To determine whether we could efficiently deliver Cas9 modRNA to hPSCs using lipofection, we synthesized Cas9-P2A-GFP modRNA containing N1-Methyl-pseudo-UTP(Hadas et al., 2019). We then transfected Cas9-P2A-GFP into undifferentiated H9 cells. We quantified GFP expression 24 hours later using flow cytometry and found we were able to achieve up to 90% transfection efficiency based on GFP positive cells (**Fig S1**). Next, to probe for the optimal amount of Cas9 modRNA and target-specific sgRNA, we made Cas9 modRNA without co-expression of GFP to KO GFP from a human embryonic stem cell (hESC) OCT4-GFP reporter line (H1 OCT4-GFP)(Zwaka and Thomson, 2003). For designing our sgRNA targeting GFP, we chose to use the GFP sgRNA sequence reported by Sanjana et al.(Sanjana et al., 2014). H1 OCT4-GFP cells were seeded in a 24-well plate and then transfected with different amounts of Cas9 modRNA and GFP sgRNA using lipofectamine Stem transfection reagent (**Fig 1A**). Four days after transfection, cells were collected to quantify the percentage of GFP-cells using flow cytometry (**Fig S2**).

**Figure 1.**
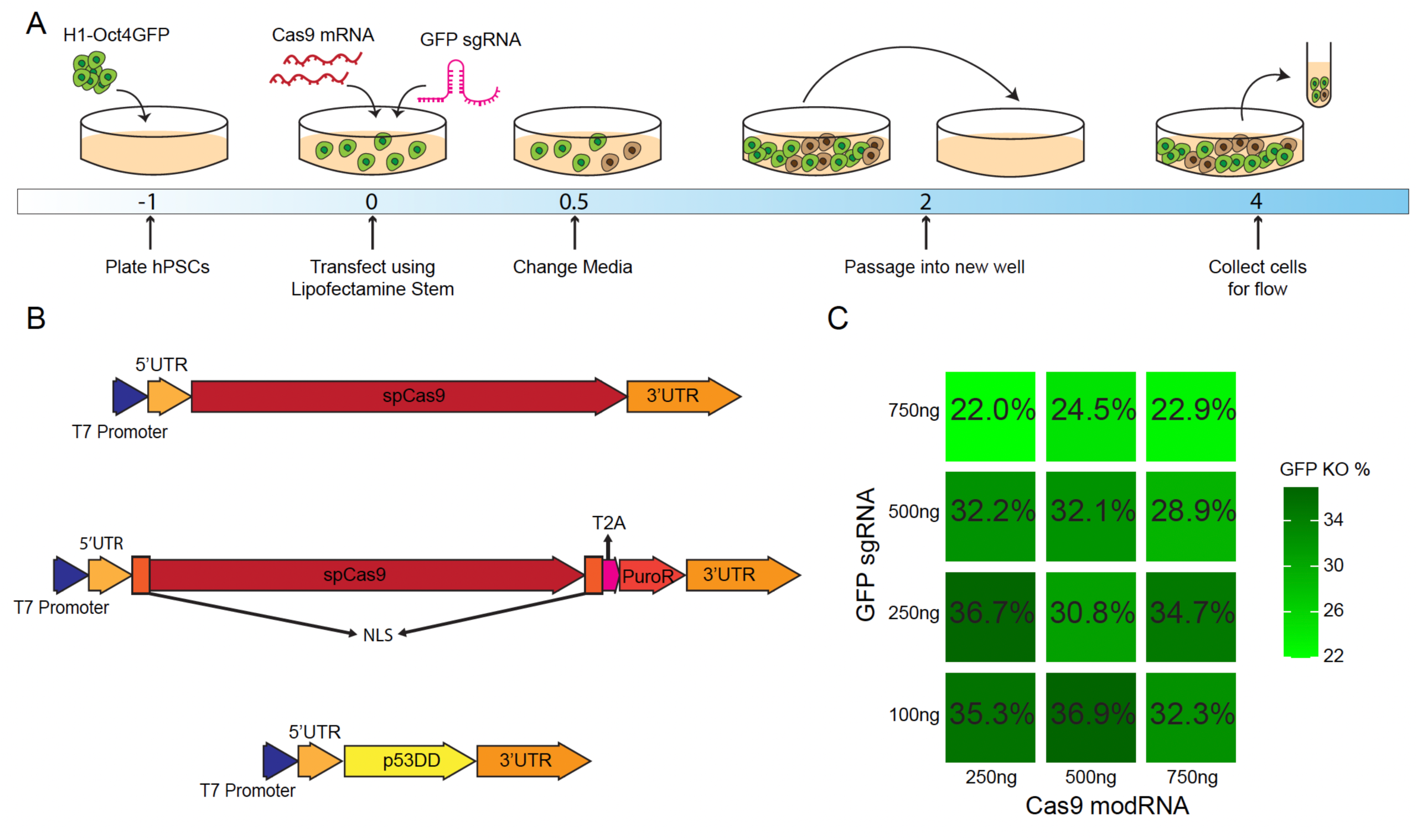
Cas9 modRNA and sgRNA efficiently knock out an integrated GFP in hPSCs. (A) Schematic diagram for knocking out GFP in H1 OCT4-GFP cells using Cas9 modRNA and *in-vitro* synthesized sgRNA. (B) DNA templates used to synthesize modRNA for Cas9, Cas9Puro, and p53DD. (C) H1 OCT4-GFP cells were cultured on iMatrix-511 in mTeSR1 and transfected with different combinations of Cas9 modRNA and GFP sgRNA. On day 4, cells were collected and GFP expression was analyzed via flow cytometry. The percentage of GFP-cells for each combination is shown in the form of a tiled heatmap.

We tested various amounts of Cas9 modRNA (250 ng, 500 ng, or 750 ng) along with different doses of GFP sgRNA (100 ng, 250 ng, 500 ng, and 750 ng) in H1 OCT4-GFP cells. We found that three Cas9+sgRNA combinations (250+100, 250+250, and 500+100) achieved the highest KO efficiency (∼36% GFP-cells on day 4) (**Fig 1B-C**). We also tested fewer amounts of Cas9 modRNA (125 ng, or 250 ng) along with fewer doses of GFP sgRNA (10 ng, 50 ng, or 100 ng), but found that these conditions performed poorly when compared to our achieved three optimal combinations (**Fig S2**). To minimize the total RNA required for transfection we decided to use the 250+100 combination for subsequent experiments.

### ModRNA-based CRISPR system efficiently generates gene KOs in hPSCs

To investigate whether our modRNA-based CRISPR system was able to efficiently knock out genes in hPSCs, we decided to target gene ***THY1*** that encodes for CD90 protein, a heavily glycosylated membrane protein that is expressed in undifferentiated hPSCs(Tang et al., 2011). We selected two potential sgRNA target sites for CD90 using ChopChop(Labun et al., 2019) (**Fig 2A**). We noticed that seeding the cells too sparsely for endogenous gene KO led to cell detachment and death. To tackle this, we decided to double our initial seeding density and include a ROCK inhibitor (Y-27632) in our culture media which led to better cell survival but reduced our transfection efficiency. We found that “CD90 sgRNA_1” was able to achieve higher KO efficiency and therefore was used for all subsequent experiments (**Fig 2B**). Next, we wanted to see if we could improve this KO efficiency for CD90 via drug selection. We synthesized Cas9-Puro modRNA, which has a puromycin resistance gene linked to the Cas9 via a P2A linker (**Fig 1B**). Due to the larger size of the Cas9Puro construct, we also tested delivery of 300 ng of Cas9Puro modRNA in addition to the previously determined 250 ng. H9 cells were seeded onto iMatrix-511 coated wells and transfected with either 250 ng of Cas9 modRNA, 250 ng of Cas9Puro, or 300 ng of Cas9-Puro modRNA. After 12 hours, cells were treated with 1 μg/ml puromycin. After 24 hours of drug selection, cells were counted using a hemocytometer. Treatment with puromycin effectively killed all cells in wells transfected with the Cas9 modRNA. However, in wells that were transfected with our Cas9Puro modRNA, we observed cell survival similar to our un-transfected, non-puro-treated control indicating our Cas9Puro modRNA could protect transfected cells from puromycin-mediated cell toxicity (**Fig 2B, C**). Additionally, we observed consistently higher cell numbers in wells that were transfected with 300 ng Cas9Puro compared to 250 ng of Cas9Puro, a difference that was statistically significant (p = 0.03, unpaired student’s t-test) **(Fig 2D)**. Due to the higher transfection efficiency using 300 ng of Cas9Puro, as indicated by higher cell survival, we used 300 ng Cas9Puro modRNA for subsequent experiments. To evaluate whether puromycin treatment increases KO efficiency, we used our Cas9Puro modRNA to knock out CD90 in H9 cells accompanied by puromycin treatment at a concentration ranging from 0 to 1 μg/ml. We observed a greater than two-fold increase in CD90 KO efficiency measured by the percentage of CD90-cells on day 5 post-transfection **(Fig 2E)**.

**Figure 2.**
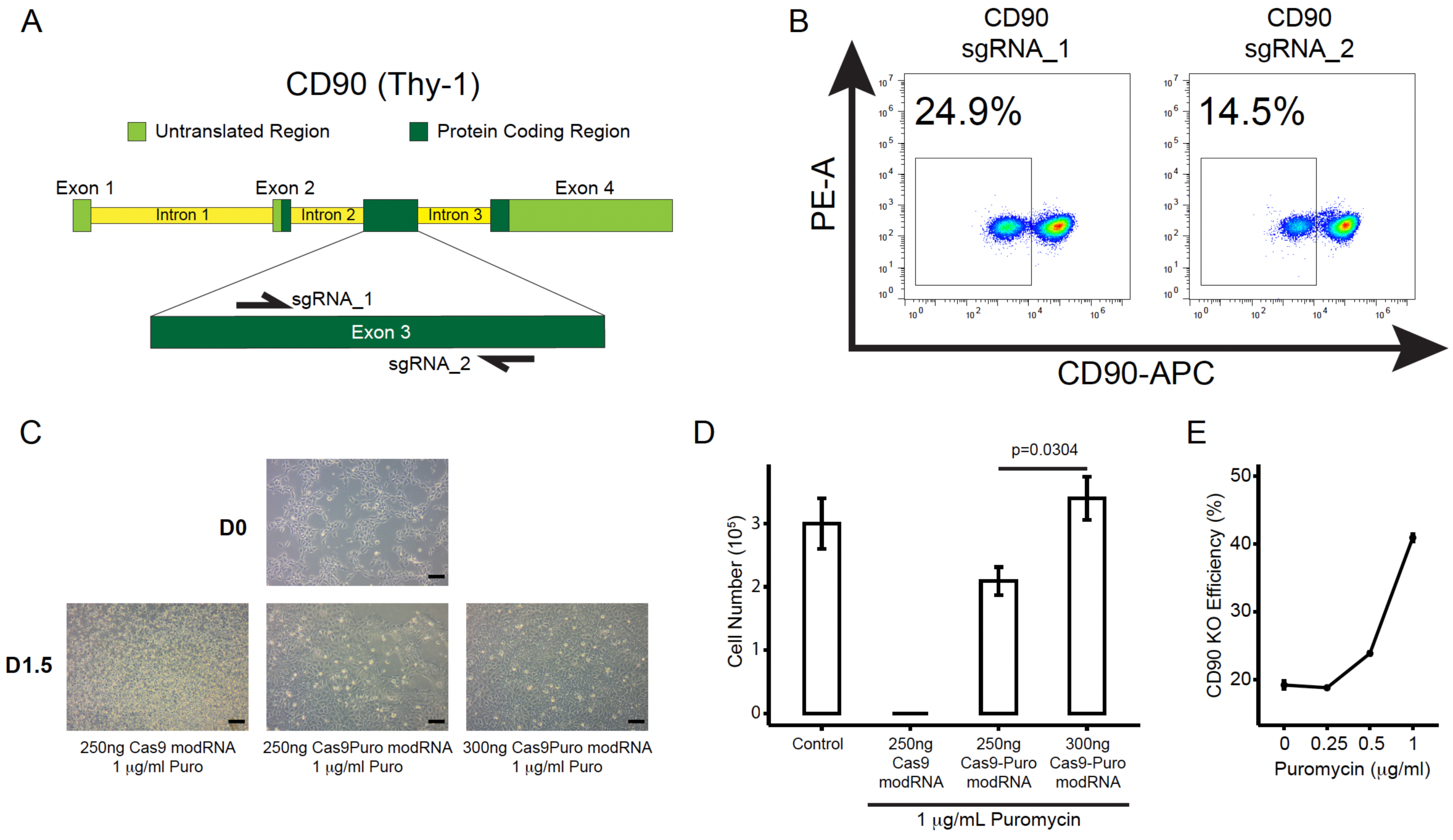
Drug Selection improved KO efficiency via Cas9Puro modRNA. (A) Schematic of sgRNA design targeting *THY1* gene, encoding CD90 protein. (B) H9 cells were cultured on iMatrix-511 in mTeSR1 and transfected with Cas9Puro modRNA and either the CD90_1 or CD90_2 sgRNA. On day 4, cells were collected and CD90 expression was analyzed via flow cytometry. Representative flow cytometry results are shown for each target design. (C) H9 cells were cultured on IMatrix-511 in mTeSR1 and transfected with either 300 ng or 250 ng of Cas9Puro modRNA or 250ng of Cas9 modRNA. Transfected cells underwent drug selection for 24 hours using puromycin beginning 12 hours after transfection. Following drug selection, cells were imaged (scale bar = 100 μm) (D) and counted using a hemocytometer (Control: n=2; 250 ng Cas9, 250ng Cas9Puro, 300ng Cas9Puro: n=3; unpaired student’s T-test). (E) H9 cells were transfected with 300 ng Cas9Puro modRNA, 100 ng CD90_1 sgRNA and underwent 24 hours of drug selection beginning 12 hours after transfection. On day 5, cells were collected and CD90 expression was analyzed by flow cytometry (n = 3).

### P53DD greatly increases modRNA-based genome editing efficiency in hPSCs

While CRISPR-Cas9 systems have been widely used to engineer genomes of a wide variety of cell types, hPSCs have proven to be exceptionally difficult to engineer due to the toxicity of DSBs in these cells. Recently, Ihry *et al*. reported that the hPSC response to Cas9 induced DSBs is mediated by p53(Ihry et al., 2018). Additionally, they showed that p53DD, a dominant negative mutant of p53, can transiently block p53 function and therefore reduce Cas9-induced toxicity in hPSCs(Ihry et al., 2018). Therefore, we decided to synthesize p53DD modRNA to use with our modRNA-based Cas9 system. We transfected H1 OCT4-GFP cells with Cas9Puro modRNA, GFP sgRNA, and p53DD modRNA in a 24-well plate **(Fig 3A)**. Wells that received p53DD modRNA had a significantly higher GFP KO efficiency measured by the percentage of GFP-cells (58.96% ± 10.17% with p53DD vs 37.13% ± 4.48% without p53DD) (**Fig 3B, C**). The difference between wells with and without p53DD was even more pronounced for CD90 KO. The KO efficiency for CD90 increased by more than two-fold with the addition of p53DD modRNA to the transfection mix (71.09% ± 9.13% vs 32.72% ± 10.83%) (**Fig 3B, C)**. Addition of p53DD modRNA to our CRISPR cocktail also abolished the need to seed cells at the higher seeding density for endogenous gene KO. To compare our modRNA-based CRISPR system to a plasmid-based system, we cloned our sgRNA target sequences (GFP and CD90) into a CRISPR plasmid containing both Cas9 and sgRNA expression cassettes(Jiang et al., 2022). Cells were then transiently transfected with either the plasmid-based CRISPR vector with corresponding gRNA sequence or our modRNA-based CRISPR cocktail. Flow cytometry of cells on day 5 demonstrated that our modRNA-based CRISPR system achieved a significantly higher KO efficiency compared to the state-of-art plasmid-based method. For CD90, while the plasmid-based method yielded 18.75% ± 5.27% KO efficiency, our modRNA-based method generated 71.09% ± 9.13% CD90 KO efficiency (p=1.56 × 10^−8^, one-way ANOVA with post-hoc Tukey’s test). (**Fig 3B, C**). Furthermore, we proved that our modRNA-based CRISPR system was also effective in editing induced pluripotent stem cells (iPSCs), with up to 63% CD90-cells on day 5 post-transfection (**Fig 3D**).

**Figure 3.**
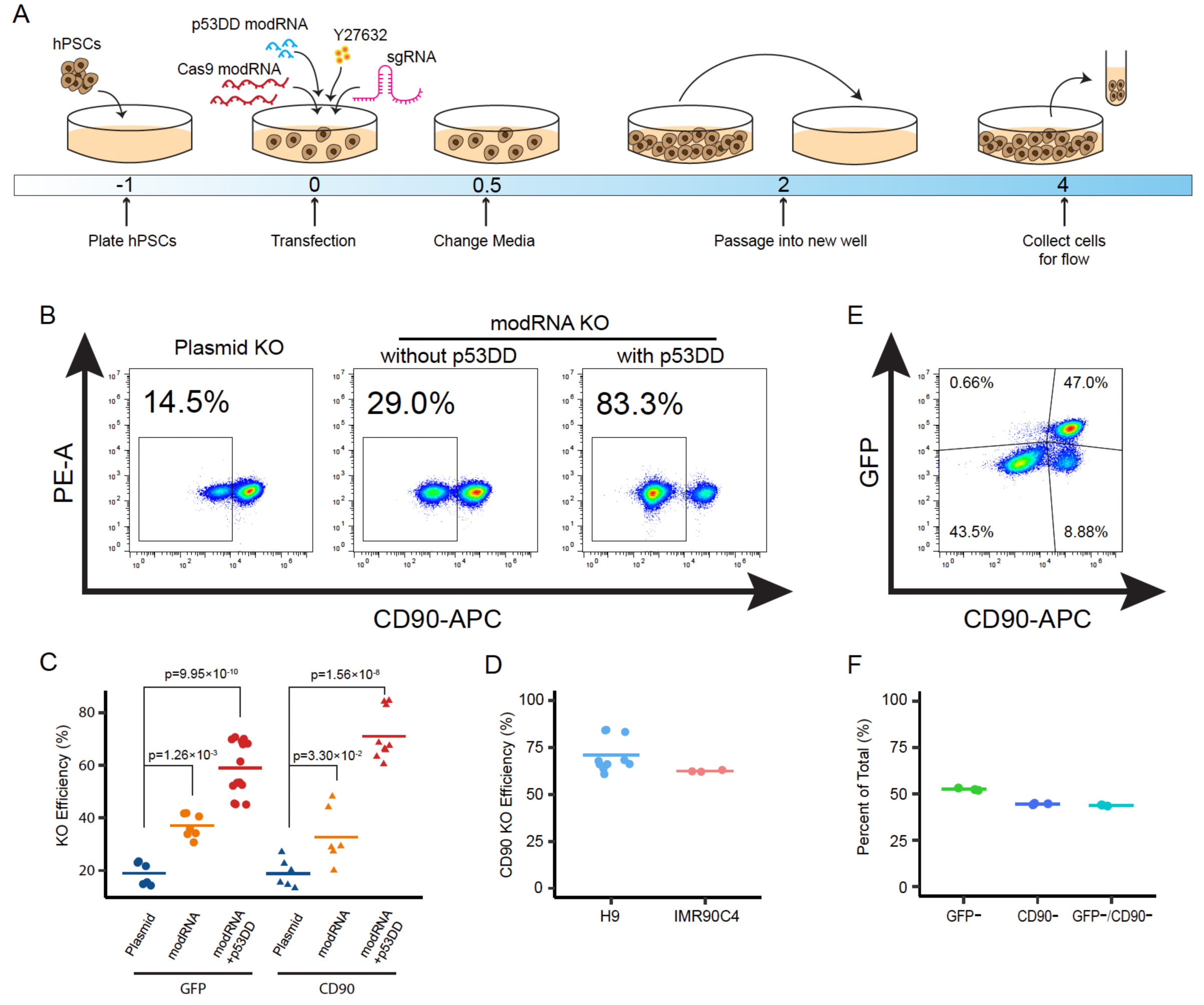
P53DD modRNA increased Cas9 modRNA mediated gene KO in hPSCs. (A) Schematic of optimal transfection protocol with the addition of p53DD modRNA. (B) H9 cells cultured on iMatrix-511 in mTeSR1 were transiently transfected with either the plasmid DNA, modRNA cocktail without or with p53DD modRNA. On day 5, cells were collected and CD90 expression was analyzed via flow cytometry. (C) Aggregated gene KO efficiencies across multiple replicates and batches for GFP and CD90, comparing results between transient plasmid DNA transfection, modRNA method without or with p53DD modRNA. (GFP/plasmid: n = 6; GFP/modRNA: n = 7; GFP/modRNA+p53DD: n = 14; CD90/plasmid: n = 6; CD90/modRNA: n = 6; CD90/modRNA+p53DD: n = 10; one-way ANOVA with post-hoc Tukey’s test). (D) IMR90C4 cells cultured on iMatrix-511 in mTeSR1 were transfected with Cas9Puro modRNA, CD90 sgRNA, and p53DD modRNA. On day 5, cells were collected and CD90 expression was analyzed via flow cytometry (n = 3). (E and F) H1 OCT4-GFP cells were cultured on iMatrix-511 in mTeSR1 using a 12-well plate, and transfected with 1200 ng Cas9Puro modRNA, 200 ng CD90_1 sgRNA, 200 ng GFP sgRNA, and 200 ng p53DD modRNA. On day 5, cells were collected and GFP/CD90 expression was analyzed via flow cytometry (n = 3). (E) Representative flow cytometry plot from day 5. (F) Quantification of flow cytometry results from day 5 cells.

Eukaryotic RNA is normally capped at the 5’-end with 7-methylguanosine (^m7^G), commonly referred to as Cap 0, and is important for translational initiation and prevents degradation of the mRNA transcript. When synthesizing modRNA, the Cap 0 structure is introduced by the addition of the anti-reverse cap analog (ARCA) to the *in vitro* transcription reaction mix. Higher order eukaryotes will instead have a Cap 1 structure, in which the first nucleotide proximal to the cap structure is methylated. Using modRNA with the Cap 1 modification can potentially abrogate the innate immune response compared to Cap 0 due to its reduced affinity for binding RIG-I, MDA5, and IFIT-1(Abbas et al., 2017; Devarkar et al., 2016; Rehwinkel and Gack, 2020; Vaidyanathan et al., 2018; Züst et al., 2011). To synthesize modRNA with the Cap 1 modification, we used site-directed mutagenesis to convert the G to an A proximal to the T7 promotor sequence in our original modRNA plasmid. Then we cloned our Cas9Puro insert into our newly synthesized modRNAc1 plasmid. In addition, we replaced the ARCA reagent with the CleanCap AG reagent. Our data showed that both cap 0 and cap 1 modRNA could efficiently knock out *THY1* (CD90) in hPSCs (**Fig S3**), indicating that the reduced immunogenicity of cap 1 modRNA did not further improve gene KO efficiency in hPSCs.

Next, we decided to examine whether our modRNA-based CRISPR system could simultaneously target multiple genomic sites and thus knock out multiple gene. We seeded our H1 OCT4-GFP cells and transfected them with Cas9Puro modRNA, GFP sgRNA, CD90 sgRNA, and p53DD modRNA. We collected cells on day 5 post-transfection to quantify GFP and CD90 expression using flow cytometry. We observed 43.5% of cells that were deficient in both GFP and CD90 expression (**Fig 3E, F**) after one single transfection.

### ABE8e modRNA outperforms its plasmid counterpart for gene KO in hPSCs

Besides CRISPR-Cas9, base editing, the introduction of single-nucleotide variants (SNVs) into the genome, is another important technique for genome editing. The adenosine deaminase base editor, ABE8e, was our base editor of choice(Richter et al., 2020). To determine if base editing efficiencies using modRNA could outperform plasmid-based delivery, we decided to knock out the *B2M* gene, a protein subunit required for surface expression of all class I major histocompatibility complex molecules via modRNA or plasmid-based ABE8e expression. Our B2M KO strategy (**Fig 4A**) was mediated by base editing the splice donor of exon 1, rendering it deactivated(Kluesner et al., 2021). Using the SpliceR program(Kluesner et al., 2021), we chose the most efficient sgRNA for B2M KO for the ABE8e system. Next, hPSCs were transfected with either plasmid or modRNA based ABE8e system. ABE8e mediated B2M KO efficiencies were measured using flow cytometry of B2M expression at day 5 post-transfection. Whereas the plasmid-based method achieved 16.33 ± 0.76% KO efficiency, our modRNA-based method generated a much higher KO efficiency (68.53 ± 3.81%) (**Fig 4B, C**). Our experiments demonstrated that our modRNA-based ABE8e system is about four times more efficient than its DNA plasmid counterpart at generating base edited splice gene KOs in hPSCs, thus illuminating the use of modRNA in this context.

**Figure 4.**
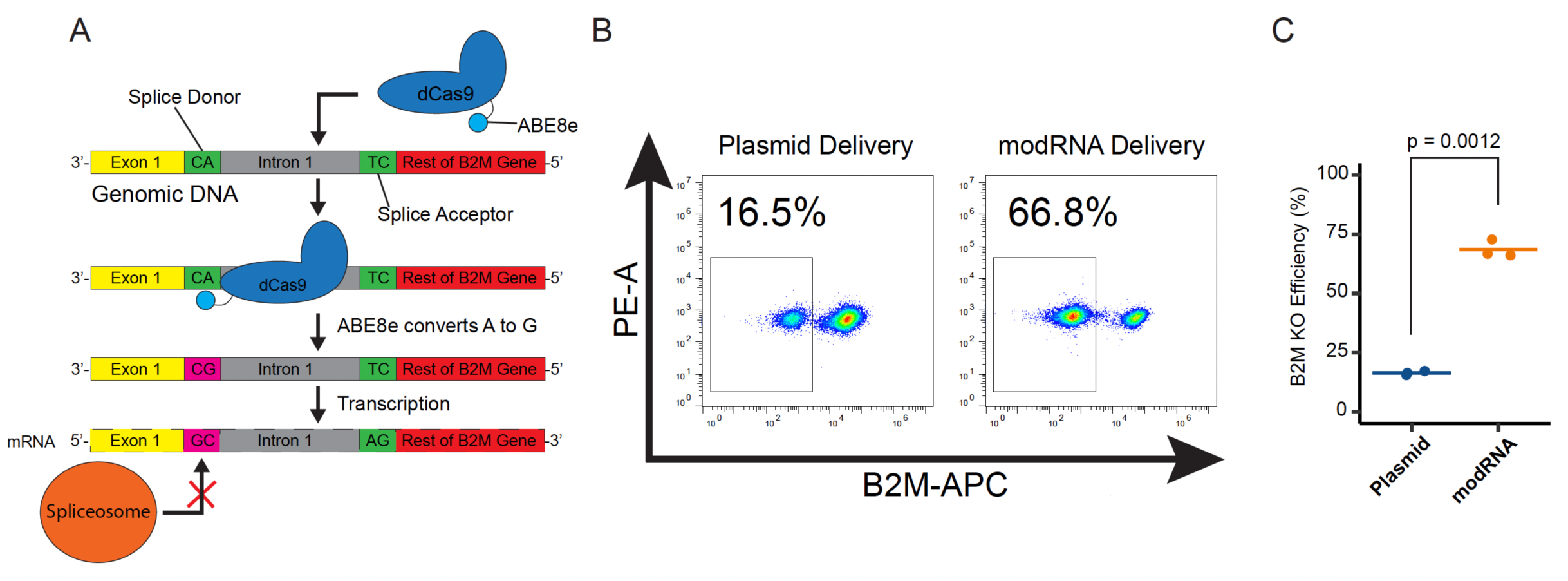
ModRNA ABE8e is more efficient over plasmid-based method. (A) Schematic of mechanism for gene KO via base editing. The dCas9 guides the fused ABE8e to the specific genomic region to perform the desired base edit. This desired base edit mutates the splice acceptor or donor region, so that after transcription, the spliceosome fails to splice out the intron or splices an exon, respectively. (B) Representative flow cytometry plots of cell population that were transfected with ABE8e + sgRNA, which were delivered in plasmid DNA or modRNA form. Cell populations were stained with a conjugated anti-B2M-APC antibody. (C) Quantification of B2M-cells following either plasmid DNA or modRNA ABE8e transfection. (p=0.0012, unpaired student’s t-test).

## DISCUSSION

Our research outlines methods for efficient CRISPR mediated gene KOs in hPSCs using a modRNA-based Cas9 or ABE8e system, which can be widely adopted for most labs without requiring electroporation or nucleofection devices. We tested the efficacy of our modRNA Cas9 system using multiple hPSC lines, including two hESC lines as well as a human iPSC line, demonstrating the general applicability. Our approach is highly flexible to a variety of experimental conditions owing to the Cas9Puro modRNA which can be used in conjunction with puromycin treatment to increase KO efficiency when high transfection efficiency is not possible for certain cell types. Integration of p53DD modRNA into our system significantly increases gene KO efficiency by reducing Cas9 induced DSB toxicity in hPSCs.

We also studied B2M gene KO in hPSCs via inactivation of the splice donor, using the ABE8e base editor. We found that modRNA ABE8e method is more efficient compared to a state-of-art plasmid format. The main advantage of using base editors for generating gene KO in hPSCs is the elimination of DSBs generated by Cas9. This abolishes the undesired chromosomal rearrangements that result from DSBs and lowers the chances of detrimental off-target insertions or deletions, thus providing a more clinically relevant genome engineering tool for hPSCs. Our modRNA ABE8e method generated a KO efficiency similar to that of our modRNA Cas9 cocktail (68.53% ± 3.81% vs. 71.09% ± 9.13%), highlighting its potential as an alternative to CRISPR-Cas9 based strategies.

In summary, we demonstrated that when CRISPR-Cas9 or ABE8e modRNA is transfected into hPSCs, it outperforms the plasmid DNA-based method. The increased efficiency of modRNA methods is likely due to lower transfection efficiency and lower Cas9 or ABE8e protein expression level in hPSCs for the plasmid-based method. By using Cas9 or ABE8e modRNA, however, it results in extremely higher transfection efficiency (more than 90%) and very high Cas9 or ABE8e expression level, ultimately generating higher KO efficiencies in hPSCs.

## Materials and Methods

### Maintenance of hPSCs

All cell culture experiments involving human pluripotent stem cell lines were approved by the Embryonic Stem Cell Oversight Committee at the Pennsylvania State University and carried out in accordance with the approved guidelines. hESCs (H1 OCT4-GFP, and H9) and human iPSCs (IMR90C4) were obtained from WiCell Research Institute (**Table S1**). hPSCs were maintained on iMatrix-511 (Iwai North America) coated plates in mTeSR1 medium (STEMCELL Technologies). Cells were regularly passaged when they reached 80-90% confluency, usually 3-4 days after the previous passage. For passaging, cell medium was aspirated and 1ml of Accutase (Innovative Cell Technologies) was added to each well. Cells were incubated at 37°C, 5% CO_2_ for 5 minutes. Dissociated cells were transferred to excess DMEM at a 1:2 (vol/vol) ratio and centrifuged at 1000 rpm for 4 minutes. New wells were precoated with 0.75 μg/ml iMatrix-511 and incubated at 37°C, 5% CO_2_ for 10 minutes. After centrifugation, cell pellet was resuspended in mTeSR1 with 5 μM Y-27632. 10,000-20,000 cells/cm^2^ were seeded onto iMatrix-511 coated wells. For regular maintenance cells were cultured in six-well plates.

### Modified mRNA (modRNA) synthesis

Cas9, Cas9Puro, Cas9GFP, p53DD, and ABE8e-GFP template DNA was PCR amplified from the donor plasmid using appropriate primers (**Table S2**). All the associated plasmids are listed in **Table S3**. The PCR product was run on a 1% Agarose gel and the band at the appropriate size was excised and the DNA extracted using the Zymoclean Gel DNA Recovery kit (Zymo Research). Purified insert DNA was cloned into the linearized modRNA plasmid (5MCS3)(K et al., 2018) using the In-Fusion Cloning Kit (Takara Bio). The DNA template for modRNA synthesis was PCR amplified from the successfully cloned 5MCS3 plasmid followed by PCR purification using DNA Clean & Concentrator-5 (Zymo Research). ModRNA was synthesized from the DNA template via *in vitro* transcription (IVT) using the MEGAscript T7 Transcription kit (ThermoFisher) supplemented with 8.1 mM ATP, 2.7 mM GTP, 8.1 mM CTP, 2.7 mM N1-methyl-pseudo-UTP, and 10 mM Anti-Reverse Cap Analog (ARCA). The IVT reaction product was treated with DNase I to remove DNA template and then purified using the MEGAclear transcription clean-up kit (ThermoFisher). RNA concentration was measured using a NanoDrop (ThermoFisher).

### sgRNA synthesis

sgRNA was synthesized using EnGen sgRNA Synthesis kit, S. pyogenes (NEB #E3322). Target specific oligos were ordered from Integrated DNA Technologies using the following template: *TTCTAATACGACTCACTATA***G**(N)_20_**<u>GTTTTAGAGCTAGA</u>**. Gene-specific target sequences for CD90 were selected using the ChopChop online tool (**Table S4**). The IVT reaction was assembled based on the manufacturer’s recommendations and the sgRNA was purified using an RNA Clean & Concentrator-5 kit (Zymo Research). RNA concentration was measured using a NanoDrop (ThermoFisher)

### Transfection of Cas9 modRNA or plasmid into hPSCs

For Cas9 mediated KO, ∼13,000 cells/cm^2^ hPSCs were seeded onto iMatrix-511 coated wells of a 24-well plate and cultured for 24 hours at 37°C, 5% CO_2_. The transfection mix was prepared using either modRNA or plasmid Cas9/Cas9-Puro, target specific sgRNA, p53DD, and lipofectamine Stem Transfection Reagent (ThermoFisher) (1:2 ratio, mass/volume) in Opti-MEM medium with GlutaMAX supplement (ThermoFisher). Before transfection, the spent medium was replaced with fresh mTeSR1 with 10 μM Y-27632. The transfection mix was incubated at room temperature for 10 minutes and then added to the well in a dropwise fashion followed by a media change 12 hours later. From then on, cells were maintained in mTeSR1 with daily media changes until cells were eventually collected for flow cytometry.

### Transfection of ABE8e modRNA or Plasmid into hPSCs

For ABE8e mediated gene KO, H9 cells were seeded onto iMatrix-511 coated wells of a 12-well plate and cultured at 37°C, 5% CO_2_. Upon reaching 30% confluency, fresh 0.5 ml mTeSR1 was added to each well, and the cells were transfected using lipofectamine Stem Transfection Reagent (ThermoFisher) in Opti-MEM medium with GlutaMAX supplement (ThermoFisher). For plasmid-based method, cells were transfected using 500 ng XLoneV3-ABE8e-GFP, 500 ng pGuide_B2M_Exon1 plasmid, and 5 μg/ml Doxycycline. For modRNA-based method, cells were transfected using 600 ng ABE8e-GFP modRNA and 200 ng B2M_Exon1_gRNA. 24 hours post transfection, a complete media change was performed using fresh mTeSR1 media, with 5 μg/mL Doxycycline supplemented to the DNA plasmid delivered wells. Cells were cultured further for another 4 days, with daily mTeSR1 media changes, and with 5 μg/mL Doxycycline for the plasmids treated cells. 5 days post-transfection, samples were analyzed for B2M expression using flow cytometry.

### Flow cytometry

hPSCs were dissociated with 1ml Accutase for 5 minutes. Cells were resuspended in FlowBuffer-1 (DPBS with 0.5% BSA) and stained with the appropriate conjugated primary antibodies (**Table S5**). Data was collected on a BD Accuri C6 Plus flow cytometer and processed using Flowjo software.

## Statistical analysis

Quantification of flow cytometry data is shown as mean ± S.D. unless otherwise stated. One-way ANOVA followed by a post-hoc Tukey’s Test was used for comparison between multiple groups. Unpaired student’s t-test was used for comparison between different experimental groups. P values ≥ 0.05 were considered not significant; P < 0.05 was considered significant.

## Author contributions

T.H., A.H., and X.L.L. designed the experiments and analyzed the results. T.H., A.H., and Y.J. performed the experiments and analyzed data. T.H. and A.H. and X.L.L. wrote the manuscript. J.Y., X.B., and X.L.L. contributed to the revision of the manuscript. X.L.L. supervised the experiments.

## Acknowledgments

This work was supported by NIH NIBIB R21EB026035 (to X.L.L.), NIH NIAMS R01 AR072731 (to J.Y.), NSF CBET-1943696 (to X.L.L.), and Penn State startup funding (to X.L.L.).

## References

Abbas, Y.M., Laudenbach, B.T., Martínez-Montero, S., Cencic, R., Habjan, M., Pichlmair, A., Damha, M.J., Pelletier, J., and Nagar, B. (2017). Structure of human IFIT1 with capped RNA reveals adaptable mRNA binding and mechanisms for sensing N1 and N2 ribose 2?-O methylations. Proc. Natl. Acad. Sci. U. S. A. 114, E2106–E2115.

Antoniou, P., Miccio, A., and Brusson, M. (2021). Base and Prime Editing Technologies for Blood Disorders. Front. Genome Ed. 3, 1.

Cong, L., Ran, F.A., Cox, D., Lin, S., Barretto, R., Habib, N., Hsu, P.D., Wu, X., Jiang, W., Marraffini, L.A., et al. (2013). Multiplex genome engineering using CRISPR/Cas systems. Science 339, 819–823.

Devarkar, S.C., Wang, C., Miller, M.T., Ramanathan, A., Jiang, F., Khan, A.G., Patel, S.S., and Marcotrigiano, J. (2016). Structural basis for m7G recognition and 2′-O-methyl discrimination in capped RNAs by the innate immune receptor RIG-I. Proc. Natl. Acad. Sci. U. S. A. 113, 596–601.

Gaudelli, N.M., Komor, A.C., Rees, H.A., Packer, M.S., Badran, A.H., Bryson, D.I., and Liu, D.R. (2017). Programmable base editing of AT to GC in genomic DNA without DNA cleavage. Nature 551, 464–471.

Hadas, Y., Sultana, N., Youssef, E., Sharkar, M.T.K., Kaur, K., Chepurko, E., and Zangi, L. (2019). Optimizing Modified mRNA In Vitro Synthesis Protocol for Heart Gene Therapy. Mol. Ther. Methods Clin. Dev. 14, 300–305.

Hsu, M.N., Chang, Y.H., Truong, V.A., Lai, P.L., Nguyen, T.K.N., and Hu, Y.C. (2019). CRISPR technologies for stem cell engineering and regenerative medicine. Biotechnol. Adv. 37, 107447.

Ihry, R.J., Worringer, K.A., Salick, M.R., Frias, E., Ho, D., Theriault, K., Kommineni, S., Chen, J., Sondey, M., Ye, C., et al. (2018). p53 inhibits CRISPR-Cas9 engineering in human pluripotent stem cells. Nat. Med. 24, 939–946.

Jiang, Y., Chen, C., Randolph, L.N., Ye, S., Zhang, X., Bao, X., and Lian, X.L. (2021). Generation of pancreatic progenitors from human pluripotent stem cells by small molecules. Stem Cell Reports 16, 2395–2409.

Jiang, Y., Hoenisch, R.C., Chang, Y., Bao, X., Cameron, C.E., and Lian, X.L. (2022). Robust genome and RNA editing via CRISPR nucleases in PiggyBac systems. Bioact. Mater.

K, S., L, T., V, B.-V., SS, D., A, K., I, S., Suknuntha, K., Tao, L., Brok-Volchanskaya, V., D’Souza, S.S., et al. (2018). Optimization of Synthetic mRNA for Highly Efficient Translation and its Application in the Generation of Endothelial and Hematopoietic Cells from Human and Primate Pluripotent Stem Cells. Stem Cell Rev. Reports 14, 525–534.

Kluesner, M.G., Lahr, W.S., Lonetree, C. lin, Smeester, B.A., Qiu, X., Slipek, N.J., Claudio Vázquez, P.N., Pitzen, S.P., Pomeroy, E.J., Vignes, M.J., et al. (2021). CRISPR-Cas9 cytidine and adenosine base editing of splice-sites mediates highly-efficient disruption of proteins in primary and immortalized cells. Nat. Commun. 12, 1–12.

Labun, K., Montague, T.G., Krause, M., Torres Cleuren, Y.N., Tjeldnes, H., and Valen, E. (2019). CHOPCHOP v3: Expanding the CRISPR web toolbox beyond genome editing. Nucleic Acids Res. 47, W171–W174.

Lian, X., Hsiao, C., Wilson, G., Zhu, K., Hazeltine, L.B., Azarin, S.M., Raval, K.K., Zhang, J., Kamp, T.J., and Palecek, S.P. (2012). Robust cardiomyocyte differentiation from human pluripotent stem cells via temporal modulation of canonical Wnt signaling. Proc. Natl. Acad. Sci. 109, E1848–E1857.

Lian, X., Zhang, J., Azarin, S.M., Zhu, K., Hazeltine, L.B., Bao, X., Hsiao, C., Kamp, T.J., and Palecek, S.P. (2013). Directed cardiomyocyte differentiation from human pluripotent stem cells by modulating Wnt/β-catenin signaling under fully defined conditions. Nat. Protoc. 8, 162–175.

Lian, X., Bao, X., Al-Ahmad, A., Liu, J., Wu, Y., Dong, W., Dunn, K.K., Shusta, E. V., and Palecek, S.P. (2014). Efficient differentiation of human pluripotent stem cells to endothelial progenitors via small-molecule activation of WNT signaling. Stem Cell Reports 3, 804–816.

Lian, X., Bao, X., Zilberter, M., Westman, M., Fisahn, A., Hsiao, C., Hazeltine, L.B., Dunn, K.K., Kamp, T.J., and Palecek, S.P. (2015). Chemically defined, albumin-free human cardiomyocyte generation. Nat Methods 12, 595–596.

Lian, X., Xu, J., Bao, X., and Randolph, L.N. (2016). Interrogating Canonical Wnt Signaling Pathway in Human Pluripotent Stem Cell Fate Decisions Using CRISPR-Cas9. Cell. Mol. Bioeng. 9, 325–334.

Liu, Z., Hui, Y., Shi, L., Chen, Z., Xu, X., Chi, L., Fan, B., Fang, Y., Liu, Y., Ma, L., et al. (2016). Efficient CRISPR/Cas9-Mediated Versatile, Predictable, and Donor-Free Gene Knockout in Human Pluripotent Stem Cells. Stem Cell Reports 7, 496–507.

Mali, P., Yang, L., Esvelt, K.M., Aach, J., Guell, M., DiCarlo, J.E., Norville, J.E., and Church, G.M. (2013). RNA-guided human genome engineering via Cas9. Science 339, 823–826.

Martin, R.M., Ikeda, K., Cromer, M.K., Uchida, N., Nishimura, T., Romano, R., Tong, A.J., Lemgart, V.T., Camarena, J., Pavel-Dinu, M., et al. (2019). Highly Efficient and Marker-free Genome Editing of Human Pluripotent Stem Cells by CRISPR-Cas9 RNP and AAV6 Donor-Mediated Homologous Recombination. Cell Stem Cell 24, 821-828.e5.

Rehwinkel, J., and Gack, M.U. (2020). RIG-I-like receptors: their regulation and roles in RNA sensing. Nat. Rev. Immunol. 20, 537–551.

Richter, M.F., Zhao, K.T., Eton, E., Lapinaite, A., Newby, G.A., Thuronyi, B.W., Wilson, C., Koblan, L.W., Zeng, J., Bauer, D.E., et al. (2020). Phage-assisted evolution of an adenine base editor with improved Cas domain compatibility and activity. Nat. Biotechnol. 2020 387 38, 883–891.

Sanjana, N.E., Shalem, O., and Zhang, F. (2014). Improved vectors and genome-wide libraries for CRISPR screening. Nat. Methods 11, 783–784.

Tang, C., Lee, A.S., Volkmer, J.P., Sahoo, D., Nag, D., Mosley, A.R., Inlay, M.A., Ardehali, R., Chavez, S.L., Pera, R.R., et al. (2011). An antibody against SSEA-5 glycan on human pluripotent stem cells enables removal of teratoma-forming cells. Nat. Biotechnol. 29, 829–834.

Vaidyanathan, S., Azizian, K.T., Haque, A.K.M.A., Henderson, J.M., Hendel, A., Shore, S., Antony, J.S., Hogrefe, R.I., Kormann, M.S.D., Porteus, M.H., et al. (2018). Uridine Depletion and Chemical Modification Increase Cas9 mRNA Activity and Reduce Immunogenicity without HPLC Purification. Mol. Ther. - Nucleic Acids 12, 530–542.

Wu, X., Dao Thi, V.L., Huang, Y., Billerbeck, E., Saha, D., Hoffmann, H.H., Wang, Y., Silva, L.A.V., Sarbanes, S., Sun, T., et al. (2018). Intrinsic Immunity Shapes Viral Resistance of Stem Cells. Cell 172, 423-438.e25.

Xu, J., Zhou, C., Foo, K.S., Yang, R., Xiao, Y., Bylund, K., Sahara, M., and Chien, K.R. (2020). Genome-wide CRISPR screen identifies ZIC2 as an essential gene that controls the cell fate of early mesodermal precursors to human heart progenitors. Stem Cells 38, 741–755.

Yilmaz, A., Peretz, M., Aharony, A., Sagi, I., and Benvenisty, N. (2018). Defining essential genes for human pluripotent stem cells by CRISPR–Cas9 screening in haploid cells. Nat. Cell Biol. 2018 205 20, 610–619.

Yu, Y., Leete, T.C., Born, D.A., Young, L., Barrera, L.A., Lee, S.J., Rees, H.A., Ciaramella, G., and Gaudelli, N.M. (2020). Cytosine base editors with minimized unguided DNA and RNA off-target events and high on-target activity. Nat. Commun. 2020 111 11, 1–10.

Züst, R., Cervantes-Barragan, L., Habjan, M., Maier, R., Neuman, B.W., Ziebuhr, J., Szretter, K.J., Baker, S.C., Barchet, W., Diamond, M.S., et al. (2011). Ribose 2’-O-methylation provides a molecular signature for the distinction of self and non-self mRNA dependent on the RNA sensor Mda5. Nat. Immunol. 12, 137–143.

Zwaka, T.P., and Thomson, J.A. (2003). Homologous recombination in human embryonic stem cells. Nat. Biotechnol. 21, 319–321.

