## Supplemental File for "Robust genome editing via modRNA-based Cas9 or base editor in human pluripotent stem cells"

^#^Co-first author

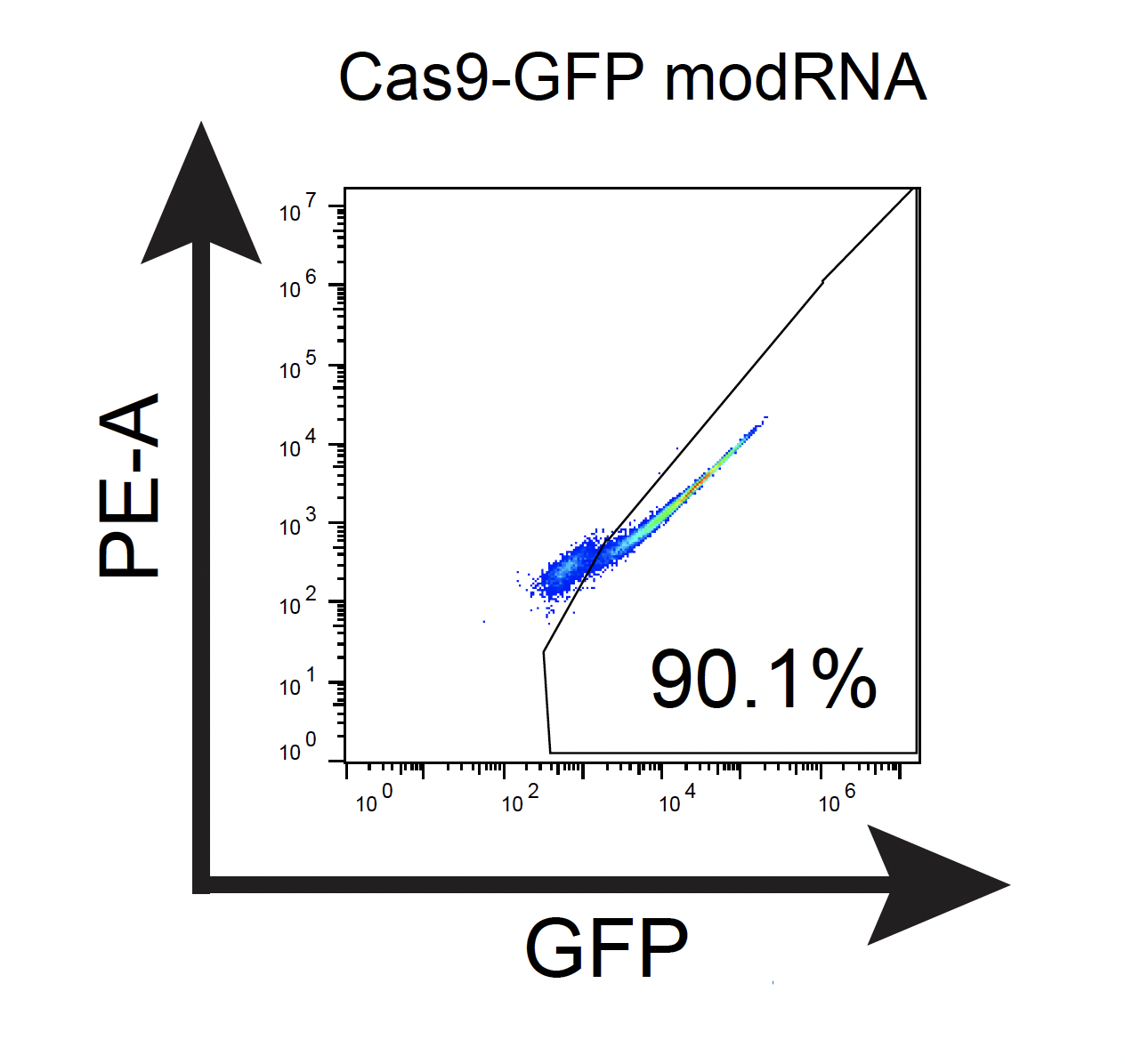

**Fig S1 Stem Transfection Reagent could efficiently deliver Cas9GFP modRNA into hPSCs.**

H9 cells were cultured on iMatrix-511 in mTeSR1 using a 24-well plate and transfected with 500 ng of Cas9GFP modRNA using lipofectamine Stem Transfection Reagent (1:2 ratio). 24 hours later GFP expression was analyzed by flow cytometry.

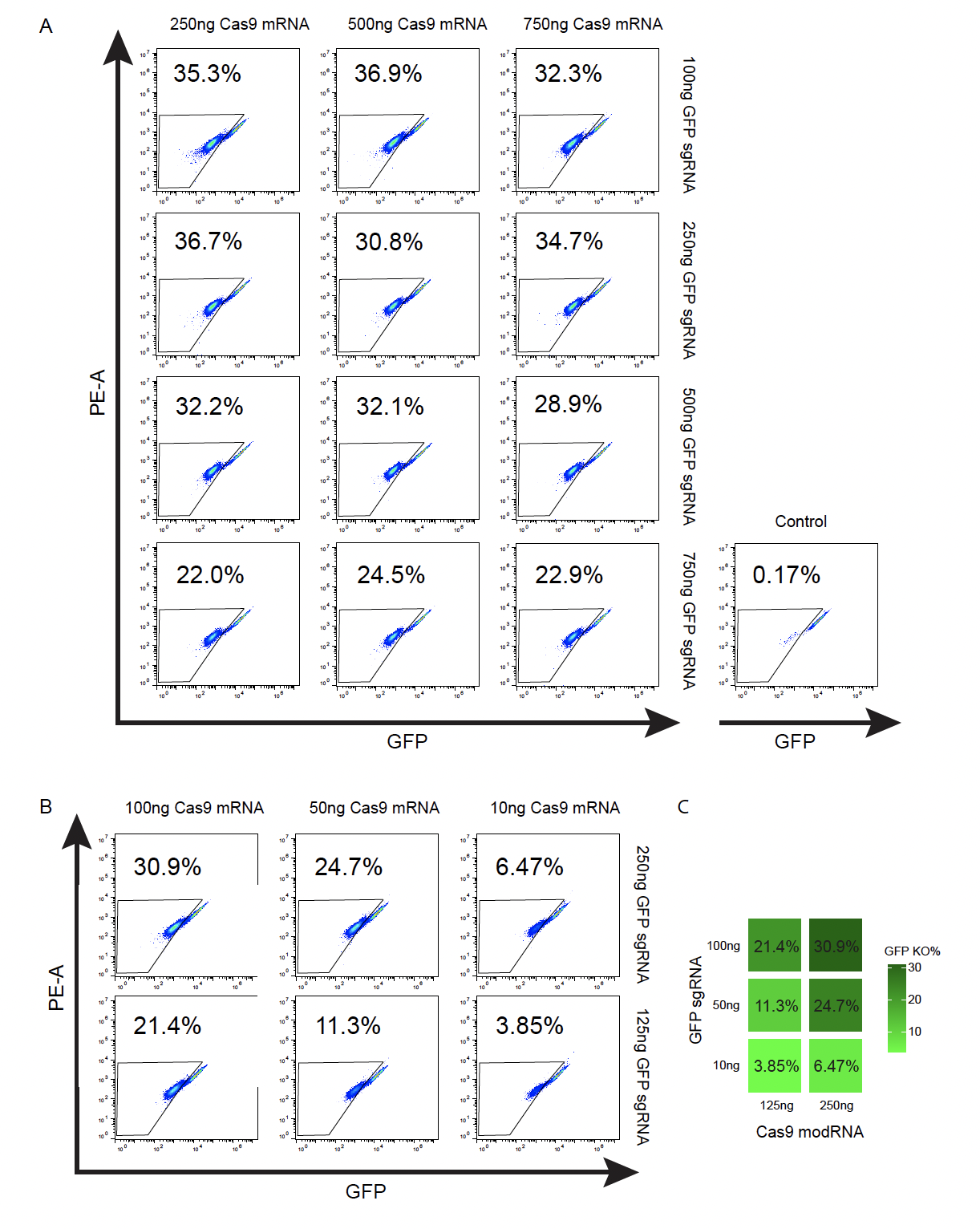

**Fig S2 Screen for optimal Cas9 modRNA and sgRNA combination by knocking out GFP in an H1 OCT4-GFP cells.** H1 OCT4-GFP cells were cultured on iMatrix-511 in mTeSR1 and transfected with different combinations of Cas9 modRNA and GFP sgRNA. On day 4, cells were collected and GFP expression was analyzed via flow cytometry. (A) Combinations of 750 ng, 500 ng, and 250 ng of Cas9 modRNA were tested with 750 ng, 500 ng, 250 ng, and 100 ng of GFP sgRNA. (B and C) Combinations of 250 ng and 125 ng of Cas9 modRNA with 100 ng, 50 ng, and 10 ng of GFP sgRNA. (B) Flow cytometry plots from day 4 cells. (C) Flow cytometry results summarized as a tiled heatmap.

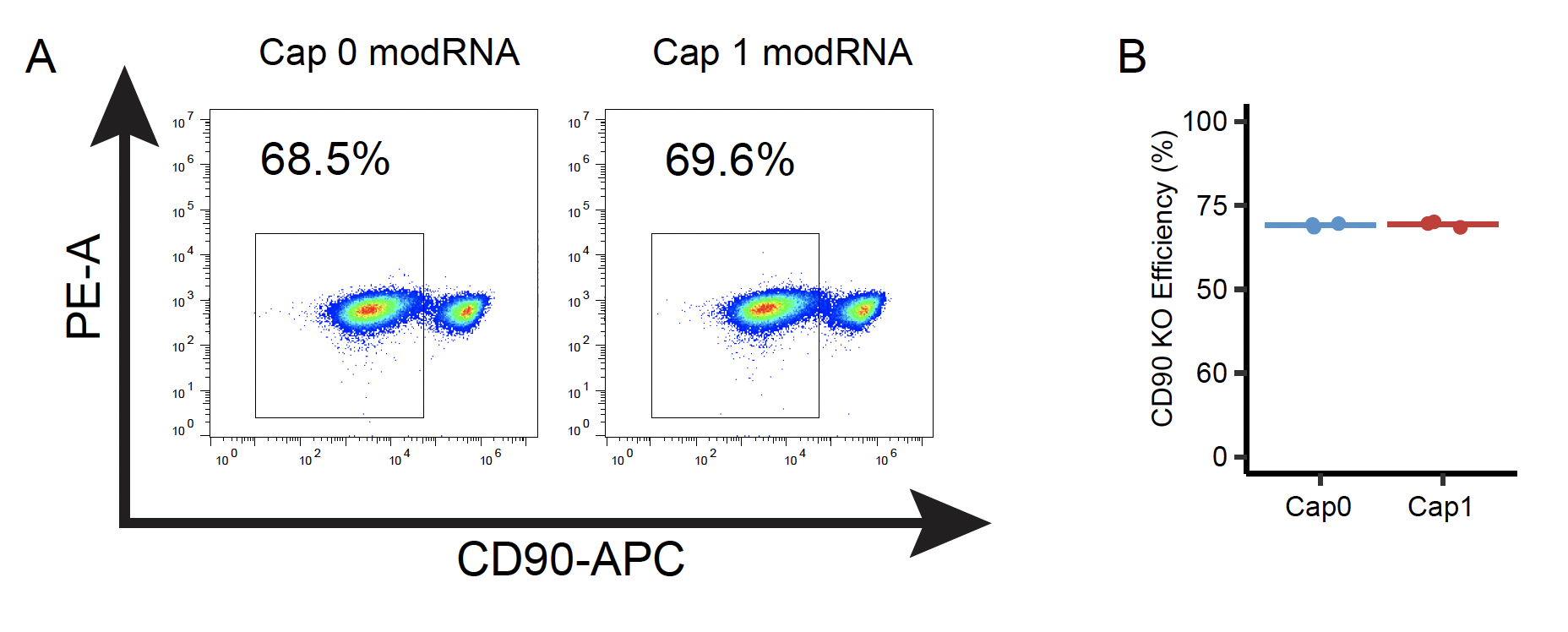

**Fig S3 Both cap 1 and cap 0 modRNA structures can mediate efficient genome editing in hPSCs.** IMR90C4 cells were cultured on iMatrix-511 in mTeSR1 using a 12-well plate and transfected with 600 ng Cas9Puro modRNA (Cap0 or Cap1), 200 ng CD90 sgRNA, and 200 ng p53DD modRNA. On day 5, cells were collected and CD90 expression was analyzed by flow cytometry. (A) Representative flow cytometry plot. (B) Quantification of flow cytometry results from day 5 cells (n = 3).

**Supplementary Table 1:** Cell lines

| **Cell Line Name** | **Source** | **Sex** | **Cell Type** |
| --- | --- | --- | --- |
| H9 | WiCell | Female | hESC |
| H1 OCT4-GFP | WiCell | Male | hESC |
| IMR90C4 | WiCell | Female | Human iPSC |

**Supplementary Table 2:** Primers used to clone target gene into 5MCS3 plasmid

| **Gene Insert** | **Sequence** |
| --- | --- |
| Cas9 | Forward: CATGGCATGCGAATTCATGGACAAGAAGTACTCCATTGGGC  Reverse: AAGCGAGCTCACTAGTTTAGTCTCCACCGAGCTGAGAG |
| Cas9-P2A-Puro | Forward: CATGGCATGCGAATTCGCCACCATGGATTACAAAGACG  Reverse: AAGCGAGCTCACTAGTTCAGGCACCGGGCTTGCG |
| Cas9-GFP | Forward: CATGGCATGCGAATTCATGGACAAGAAGTACTCCATTGGGC  Reverse: AAGCGAGCTCACTAGTTTACTTGTACAGCTCGTCCATGCC |
| p53DD | Forward: CATGGCATGCGAATTCGCCACCATGACTGCCATGG  Reverse: AAGCGAGCTCACTAGTTCAGTCTGAGTCAGGCCCC |
| ABE8e-GFP | Forward: CATGGCATGCGAATTCGCCACCATGAAACGGACAGC  Reverse: AAGCGAGCTCACTAGTTTATACCTTACGCTTCTTCTTTGGC |

**Supplementary Table 3:** Plasmids

| **Plasmid** | **Addgene #** |
| --- | --- |
| 5MCS3 | Not Available |
| XLoneV3-ABE8e-GFP | Pending submission |
| 5MCS3c0-ABE8e-P2A-GFP | Addgene, #178177 |
| pCE-mp53DD | Addgene, #41856 |
| pGuide (for DNA plasmid gRNA delivery) | Addgene, #64711 |
| modRNAc0-Cas9 | Addgene, #170180 |
| modRNAc0-Cas9-2A-GFP | Addgene, #170181 |
| modRNAc0-Cas9-2A-Puro | Addgene, #172855 |
| modRNAc0-p53DD | Addgene, #176902 |
| PB-CRISPR | Addgene, #160047 |

**Supplementary Table 4:** gRNA Sequences

| **Targeting Gene** | **Target Sequence** |
| --- | --- |
| ABE8e B2M Exon 1 Splice Donor | ACTCACGCTGGATAGCCTCC |
| GFP | GGGCGAGGAGCTGTTCACCG |
| CD90_1 | CATGGCGGCAGTCCAGACGA |
| CD90_2 | GCCTTCACTAGCAAGGACGA |

**Supplementary Table 2:** Antibodies

| **Antibody** | **Source/Host and Isotype/Clone/Catalog#** | **Application** |
| --- | --- | --- |
| B2M-APC | Biolegend/ mouse IgG_1_ / Clone:2M2 / 316312 | 1:400 (FC) |
| CD90-APC | Biolegend/ mouse IgG_1_/ Clone:5E10/ 328113 | 1:60 (FC) |
